# Mice Humanized for Major Histocompatibility Complex and Angiotensin-Converting Enzyme 2 with High Permissiveness to SARS-CoV-2 Omicron Replication

**DOI:** 10.1101/2022.12.01.518541

**Authors:** Fabien Le Chevalier, Pierre Authié, Sébastien Chardenoux, Maryline Bourgine, Benjamin Vesin, Delphine Cussigh, Yohann Sassier, Ingrid Fert, Amandine Noirat, Kirill Nemirov, François Anna, Marion Bérard, Françoise Guinet, David Hardy, Pierre Charneau, François Lemonnier, Francina Langa-Vives, Laleh Majlessi

## Abstract

Human Angiotensin-Converting Enzyme 2 (hACE2) is the major receptor enabling host cell invasion by SARS-CoV-2 via interaction with Spike glycoprotein. The murine ACE2 ortholog does not interact efficiently with SARS-CoV-2 Spike and therefore the conventional laboratory mouse strains are not permissive to SARS-CoV-2 replication. Here, we generated new *hACE2* transgenic mice, which harbor the *hACE2* gene under the human keratin 18 promoter, in C57BL/6 “HHD-DR1” background. HHD-DR1 mice are fully devoid of murine Major Histocompatibility Complex (MHC) molecules of class-I and -II and express only MHC molecules from Human Leukocyte Antigen (HLA) HLA 02.01, DRA01.01, DRB1.01.01 alleles, widely expressed in human populations. We selected three transgenic strains, with various *hACE2* mRNA expression levels and distinctive profiles of lung and/or brain permissiveness to SARS-CoV-2 replication. Compared to the previously available B6.K18-ACE2^2Prlmn/JAX^ mice, which have limited permissiveness to SARS-CoV-2 Omicron replication, these three new hACE2 transgenic strains display higher levels of *hACE2* mRNA expression, associated with high permissiveness to the replication of SARS-CoV-2 Omicron sub-variants. As a first application, one of these MHC- and ACE2-humanized strains was successfully used to show the efficacy of a lentiviral vector-based COVID-19 vaccine candidate.

## Introduction

With the persistence of the COVID-19 pandemic, the research for second-generation vaccines and drugs remains an important public health issue worldwide and therefore new research tools are still indispensable. Human Angiotensin-Converting Enzyme 2 (hACE2) is the major receptor enabling host cell invasion by SARS-CoV-2 via interaction with Spike glycoprotein (Hoffmann *et al*, 2020). The murine ACE2 ortholog does not interact properly with Spike and therefore the conventional laboratory mouse strains are not permissive to SARS-CoV-2 replication. C57BL/6 mice transgenic for *hACE2* (B6.K18-ACE2^2Prlmn/JAX^) (McCray *et al*, 2007) are available at JAX Laboratories. We also generated a new C57BL/6 transgenic strain carrying *hACE2* gene under the human keratin 18 (K18) promoter, namely “B6.K18-hACE2^IP-THV^”. Compared to B6.K18-ACE2^2Prlmn/JAX^ mice, the B6.K18-hACE2^IP-THV^ strain has distinctive characteristics, including higher *hACE2* mRNA expression and strong permissiveness to viral replication in the brain, in addition to the lung (Ku *et al*, 2021a; Vesin *et al*, 2022). In the present study, we generated three new K18-hACE2 transgenic mice in the “HHD-DR1” genetic background. HHD-DR1 mice are fully devoid of murine Major Histocompatibility Complex (MHC) molecules of class-I (MHC-I) and -II (MHC-II) and express only MHC-I and -II molecules from Human Leukocyte Antigen (HLA) alleles, i.e., HLA 02.01, DRA01.01, DRB1.01.01, widely expressed in human populations, notably in Caucasian, Asian and African populations. More precisely, HLA 02.01 is expressed by 50% of Caucasian, 50% of Asian and 35% of African population, while DR1 is expressed by 20% of Caucasian, 10% of Asian and 8% of African populations. Therefore, in the HHD-DR1 hACE2 transgenic mice, antigen presentation and cognate interactions between MHC-I and -II molecules and repertoire of T-Cell Receptors for antigen (TCR) reproduce key molecular aspects of antigen presentation in humans.

HHD-DR1 mice are triple transgenic for HLA 02.01, DRA01.01, DRB1.01.01 and quintuple knocked-out for murine β2-microglobulin (β2m), H-2D^b^, I-A_α_^b^, I-A_β_^b^ and I-E_β_^b^. Now homozygote for all these loci, HHD-DR1 mice resulted from: (i) an initial crossing of H2D^b^ KO mice with mice *Cre Lox*-deleted of a complete 80 kb-long genomic region, encompassing the genes encoding I-A_α_^b^, I-A_β_^b^, I-E_β_^b^ MHC-II molecules (Madsen *et al*, 1999) and selection, to identify mice having linked by crossing over the MHC deleted region and the H-2D^b^ KO locus, (ii) a subsequent crossing to the first generation HHD mice, transgenic for a fusion of HLA A2.01 and human β2m single chain and double KO for murine β2m and H-2D^b^ (Pascolo *et al*, 1997), followed by (iii) a final crossing to another HHD strain, double transgenic for HLA DRA01.01, DRB1.01.01 mice (Altmann *et al*, 1995; Pajot *et al*, 2004). Therefore, any possibility of epitope presentation by the H-2D^b^ heavy chain alone (Allen *et al*, 1986; Potter *et al*, 1984; Potter *et al*, 1985) or by a putative HLA DRA.01.01 - I-E_β_^b^ hybrid MHC-II molecule is fully discarded in HHD-DR1 mice (Bix & Raulet, 1992; Lawrance *et al*, 1989).

The HHD-DR1 *hACE2* transgenic strains generated in the present study are permissive to SARS-CoV-2 replication, either in the lung or in the lung and brain. Compared to B6.K18-ACE2^2Prlmn/JAX^ transgenic mice (McCray *et al*., 2007), which have weak permissiveness to SARS-CoV-2 Omicron replication (Halfmann *et al*, 2022), these HHD-DR1 hACE2 transgenic strains express higher amounts of *hACE2* mRNA, comparable to those detected in the B6.K18-hACE2^IP-THV^ strain (Ku *et al*., 2021a), which is correlated with their marked permissiveness to SARS-CoV-2 Omicron replication, in addition to previous SARS-CoV-2 variants. As a first application, in one of the established MHC- and ACE2-humanized strains, we demonstrated the strong efficacy of a lentiviral vector-based COVID-19 vaccine candidate, the mode of action of which is largely dependent on T-cell immunity (Ku *et al*., 2021a; Ku *et al*, 2021b; Majlessi & Charneau, 2021; Vesin *et al*., 2022).

## Results

### *hACE-2* transgenesis in HHD-DR1 mice

Two technologically distinct transgenesis methods were performed by: (i) pronuclear DNA micro-injection of a 6.8-kb Hpal I-XbaI fragment from the pK18-hACE2 vector (McCray *et al*., 2007) into the fertilized HHD-DR1 eggs, or (ii) sub-zonal micro-injection of the integrative LV::pK18-hACE2 lentiviral vector under pellucida of fertilized HHD-DR1 eggs. With the former approach, usually multiple head-to-tail tandem copies of the transgene are inserted into a unique site of the host genome (Haruyama *et al*, 2009), whereas with the latter technology, the transgene – not in tandem – can be integrated at numerous potential integration sites (Dussaud *et al*, 2018).

The progenies were studied for integration of *hACE2* gene. The *hACE2*^+^ F0#8 male, F0#9 female and F0#11 female founders were identified in the DNA fragment-based transgenesis. The *hACE2*^+^ F30# male and F0#47 male founders were identified in the lentiviral vector-based transgenesis (Table S1). To stabilize the progenies, each founder was backcrossed to HHD-DR1 non-*hACE2* transgenic mice to preserve the homozygous status for: (i) HLA 02.01, DRA01.01 and DRB1.01.01 humanized loci, and (ii) KO genes encoding for murine β2m, H-2D^b^, I-A_α_^b^, I-A_β_^b^ and I-E_α_^b^.

### Permissiveness of the *hACE2*^+^ offspring to SARS-CoV-2 replication

First, the *hACE2*^*+*^ offspring of the founders generated by the DNA fragment-based transgenesis were evaluated for their permissiveness to SARS-CoV-2 replication. Individuals from the previously described B6.K18-hACE2^IP-THV^ strain (Ku *et al*., 2021a; Vesin *et al*., 2022) were also included as positive controls. Mice were inoculated intranasally (i.n.) with 0.3 × 10^5^ TCID_50_ of a Delta SARS-CoV-2 clinical isolate (Planas *et al*, 2021). Four days post-inoculation (dpi), viral loads were determined in the lung and brain by E_CoV-2_ (E)-or sub-genomic E (Esg)-specific quantitative real-time (qRT)-PCR. The latter measures only active viral replication (Chandrashekar *et al*, 2020b; Tostanoski *et al*, 2020; Wolfel *et al*, 2020).

The progeny of the founder F0#8 was not permissive to SARS-CoV-2 replication (Fig 1A). In contrast, the progeny of the founder F0#9, hereafter named “HHD-DR1.ACE2^Hu1^”, displayed a notable lung – but not brain – permissiveness to SARS-CoV-2 replication. The progeny of the founder F0#11, hereafter named “HHD-DR1.ACE2^Hu2^”, was not only markedly permissive to SARS-CoV-2 replication in their lung (Fig 1A, top) but also displayed significant Esg RNA contents in their brain (Fig 1A, bottom). The viral RNA contents in the lungs of HHD-DR1.ACE2^Hu1^ and HHD-DR1.ACE2^Hu2^ mice were statistically comparable to those of the previously described B6.K18-hACE2^IP-THV^ strain. The main characteristics of these strains are recapitulated in the Table S1. Comparative assessment of *hACE2* mRNA transcription showed that the non-permissive F0#8 offspring had detectable but the lowest *hACE2* mRNA expression level in the lung and brain (Fig 1B). Such transcription levels were obviously not sufficient to allow productive SARS-CoV-2 infection. Analysis of the *hACE2* mRNA transcription levels in both lung and brain established the following hierarchy: B6.K18-hACE2^IP-THV^ = HHD-DR1.ACE2^Hu2^ > HHD-DR1.ACE2^Hu1^ > F0#8 offspring (Fig 1B).

**Figure 1.**
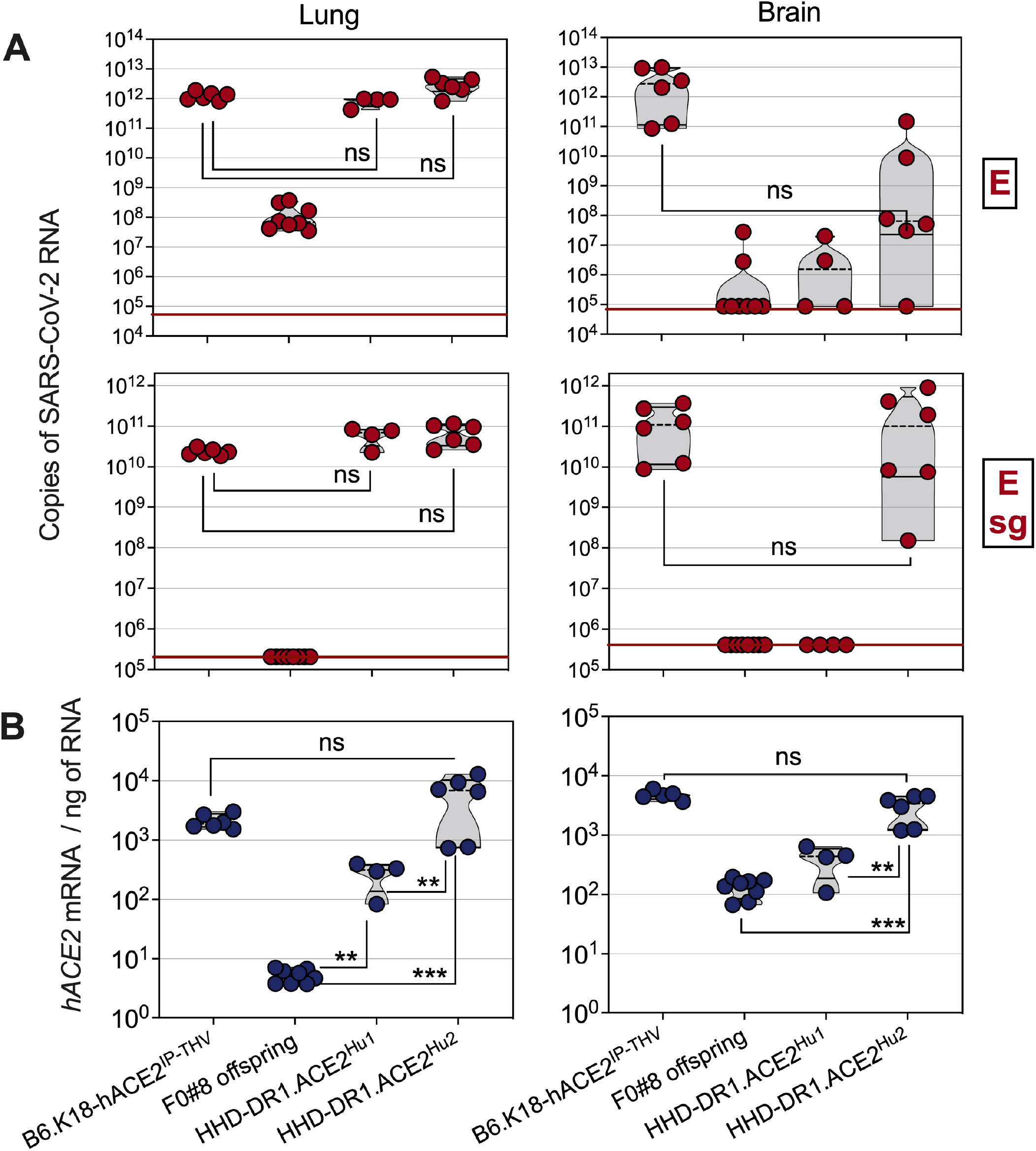
Permissiveness of HHD-DR1.ACE2^Hu1^ and HHD-DR1.ACE2^Hu2^ transgenic mice to SARS-CoV-2 replication. *hACE2*^*+*^ offspring from the founders #8, #9 (HHD-DR1.ACE2^Hu1^) or #11 (HHD-DR1.ACE2^Hu2^), resulting from a conventional DNA-based transgenesis, or B6.K18-hACE2^IP-THV^ positive control mice were inoculated i.n. with 0.3 × 10^5^ TCID_50_/mouse of SARS-CoV-2 Delta variant. **A** Lung and brain viral RNA contents were determined by E-(top) or sub-genomic Esg-(bottom) specific qRT-PCR at 4 dpi. Red lines indicate the qRT-PCR detection limits. **B** Quantification of *hACE-2* mRNA in the lung and brain of the same mice. Statistical significance was evaluated by Mann-Whitney test (*= *p* < 0.05, **= *p* < 0.01, ***= *p* < 0.001).

In the lentiviral vector-based transgenesis, the progeny of the founder F0#30 was not permissive to SARS-CoV-2 Delta replication, whereas the progeny of the founder F0#47, hereafter named “HHD-DR1.ACE2^Hu3^”, displayed strong lung (Fig 2A) and brain (Fig 2B) permissiveness to SARS-CoV-2 Delta replication. Comparative assessment of the *hACE2* mRNA transcription levels in the lung (Fig 2C) and brain (Fig 2D) of the F0#30 offspring and HHD-DR1.ACE2^Hu3^ mice established a positive correlation between the *hACE2* transgene expression levels and the permissiveness to viral replication.

**Figure 2.**
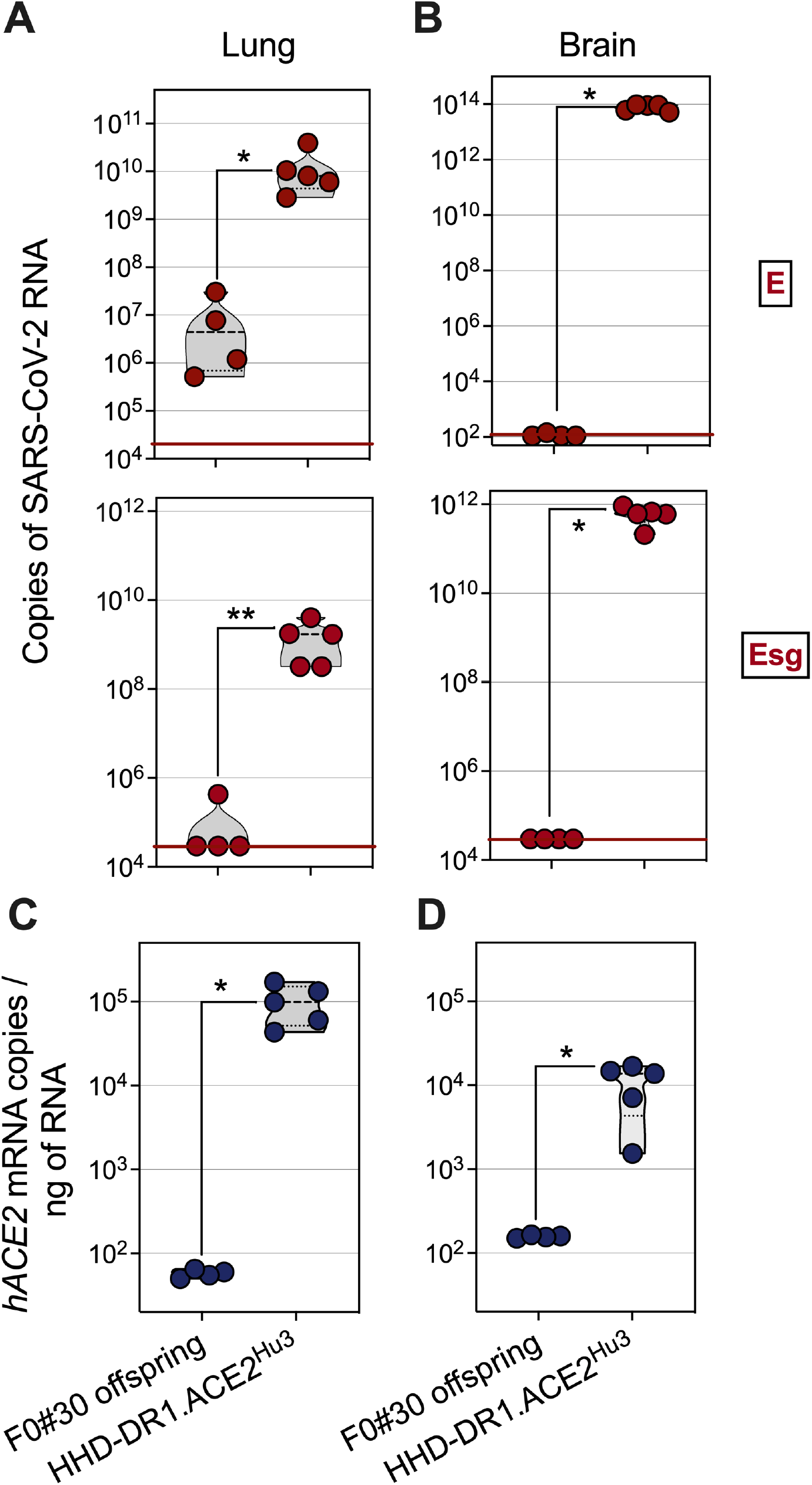
Permissiveness of HHD-DR1.ACE2^Hu3^ transgenic mice to SARS-CoV-2 replication. *hACE2*^*+*^ offspring from the founders #30 or #47 (HHD-DR1.ACE2^Hu3^), resulting from a lentiviral vector-based transgenesis, were inoculated i.n. with 0.3 × 10^5^ TCID_50_/mouse of SARS-CoV-2 Delta variant. **A-B** Lung **(A)** and brain **(B)** viral RNA contents were determined by E-specific (top) or sub-genomic Esg-specific (bottom) qRT-PCR at 4 dpi. **C-D** Quantification of *hACE-2* mRNA in the lung and brain of the same mice. Statistical significance was evaluated by Mann-Whitney test (*= *p* < 0.05, **= *p* < 0.01).

Therefore, based on the transgenesis method, *hACE2* expression levels and permissiveness of the organs to SARS-CoV-2 replication, we defined three distinct humanized HHD-DR1.ACE2^Hu^ strains (Table S1).

### SARS-CoV-2-related brain inflammation in HHD-DR1.ACE2^Hu3^ mice

HHD-DR1.ACE2^Hu3^ mice, which had the highest viral loads notably in the brain, were further characterized for their infection-mediated inflammation after inoculation with SARS-CoV-2 Delta. At 4 dpi, as evaluated by qRT-PCR study of 20 pro-or anti-inflammatory analytes, applied to RNA extracted from total lung homogenate, no sizable modification of the inflammatory transcriptome was observed in the lungs of the permissive HHD-DR1.ACE2^Hu3^ mice, compared to the non-permissive F0#30 offspring (Fig 3A). In net contrast, at this time point, the brain of the permissive HHD-DR1.ACE2^Hu3^ mice displayed a marked inflammatory status, compared to the non-permissive F0#30 offspring (Fig 3B). This brain inflammation was characterized by statistically significant upregulation of IFN-α, IFN-γ, TNF-α, IL-1β, IL-5, IL-6, IL-12p40, CCL2, CCL3, CCL5, CXCL9 and CXCL10 (Fig 3B), in a correlative manner (Fig 3C). No correlation was observed between IL-2 and IL-4 expression and the former.

**Figure 3.**
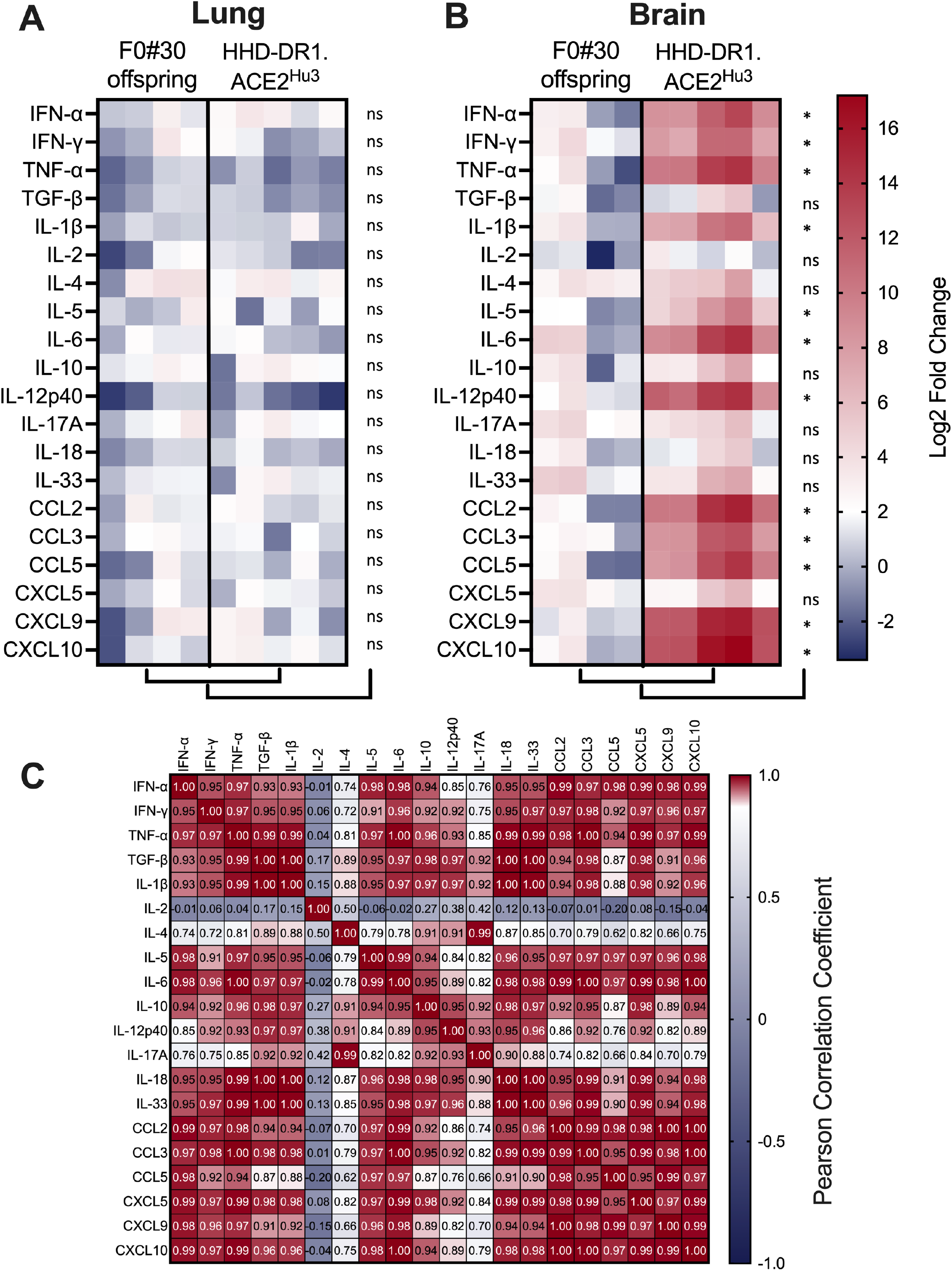
Inflammatory status of lungs and brain of HHD-DR1.ACE2^Hu3^ transgenic mice after SARS-CoV-2 inoculation. **A, B** Heatmaps represent log_2_ fold change in cytokine and chemokine mRNA expression in the lungs **(A)** or brain **(B)** of F0#30 offspring or HHD-DR1.ACE2Hu3 transgenic mice at 4 dpi (*n* = 4-5/group) after inoculation of SARS-CoV-2 Delta variant. Mice are those shown in the Figure 2. Data were normalized versus untreated controls. Statistical significance was evaluated by Mann-Whitney test (*= *p* < 0.05, ns = not significant). **C** Pearson correlation coefficient of the analytes studied in the brain of SARS-CoV-2 Delta-inoculated HHD-DR1.ACE2Hu3 mice.

### Permissiveness of HHD-DR1.ACE2 mice to replication of SARS-CoV-2 Omicron sub-variants

The previously available B6.K18-ACE2^2Prlmn/JAX^ transgenic mice are much less permissive to Omicron replication than to the other previously emerged SARS-CoV-2 variants (Halfmann *et al*., 2022). In net contrast, the B6.K18-hACE2^IP-THV^ mice, that we previously generated (Ku *et al*., 2021a), allow the replication of Omicron as efficiently as other SARS-CoV-2 variants like Delta (Vesin *et al*., 2022). Correlatively, B6.K18-^hACE2IP-THV^ mice express significantly higher amounts of *hACE2* mRNA in lungs and brain than B6.K18-ACE2^2Prlmn/JAX^ (Ku *et al*., 2021a).

Here, we first evaluated the permissiveness of the three HHD-DR1.ACE2^Hu1/2/3^ lineages to SARS-CoV-2 Omicron BA.1 sub-variant. As determined at 4 dpi, by SARS-CoV-2 E-or Esg-specific qRT-PCR, the lungs of all three HHD-DR1.ACE2^Hu1/2/3^ strains were strongly permissive to the replication of Omicron BA.1 (Fig 4A). The brain viral RNA content of HHD-DR1.ACE2^Hu1^ was not studied here, as we previously determined the brain unpermissiveness in this strain to SARS-CoV-2 replication (Fig 1A). In HHD-DR1.ACE2^Hu2^ mice, the brain E viral RNA content was relatively weak and no brain Esg viral RNA was detected (Fig 4B). In contrast, substantial Omicron BA.1 replication was detected in the HHD-DR1.ACE2^Hu3^ mice which, like the previously described B6.K18-hACE2^IP-THV^ mice, resulted from lentiviral vector-based transgenesis (Fig 4B).

**Figure 4.**
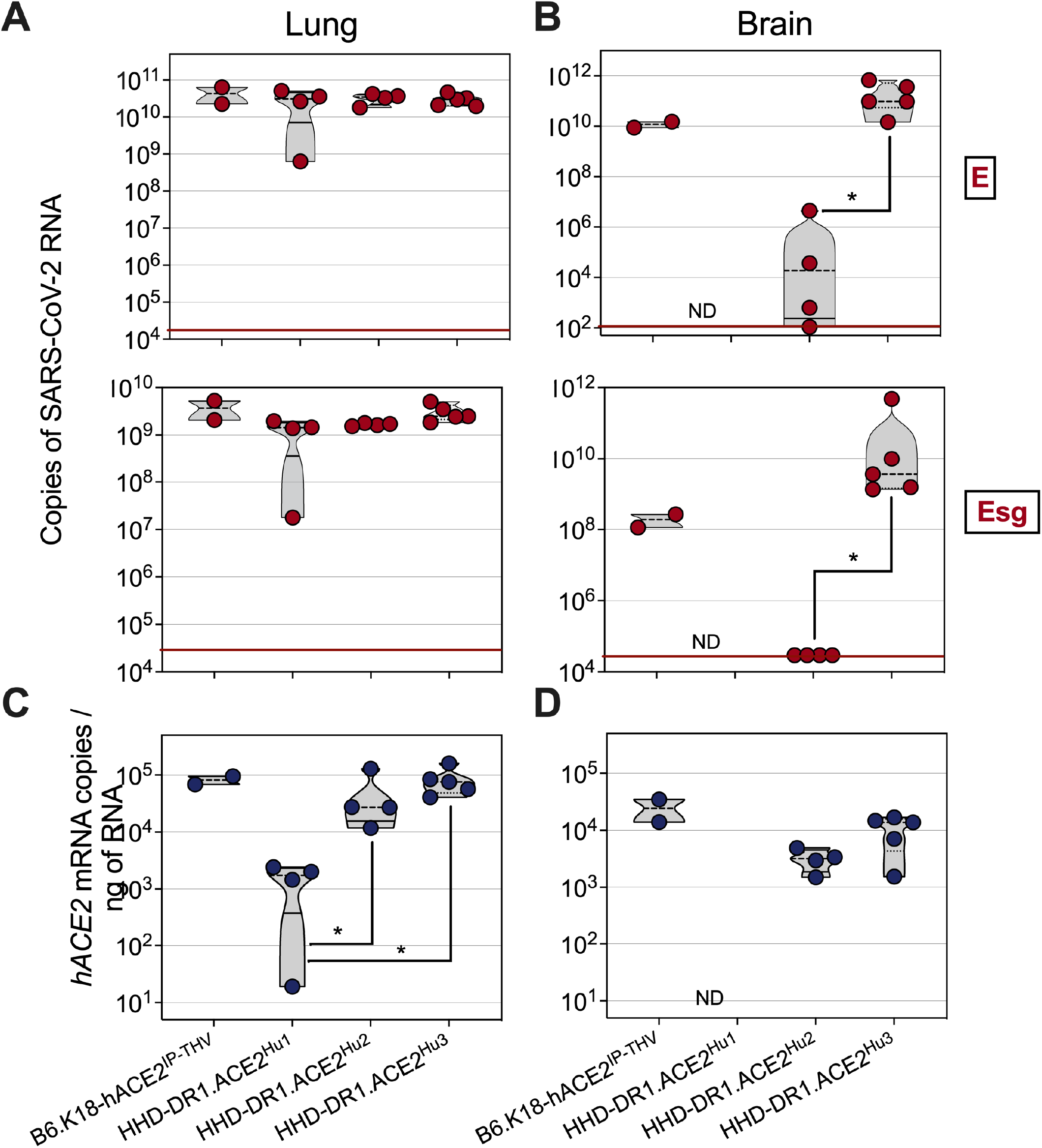
Permissiveness of HHD-DR1.ACE2Hu1/2/3 mice to replication of SARS-CoV-2 Omicron BA.1 sub-variant. HHD-DR1.ACE2Hu1/2/3 mice or B6.K18-hACE2IP-THV positive control mice were inoculated i.n. with 0.3 × 10^5^ TCID50/mouse of SARS-CoV-2 Omicron BA.1. **A-B** Lung **(A)** and brain **(B)** viral RNA contents were determined by E-specific (top) or sub-genomic Esg-specific (bottom) qRT-PCR at 4 dpi. Statistical significance was evaluated by Mann-Whitney test (*= *p* < 0.05). **C-D** Comparative quantification of *hACE-2* mRNA in the lung **(C)** and brain **(D)** of individual B6.K18-hACE2IP-THV or HHD-DR1.ACE2Hu1/2/3 transgenic mice.

Analysis of *hACE2* mRNA expression levels established the following hierarchy: HHD-DR1.ACE2^Hu3^ = HHD-DR1.ACE2^Hu2^ > HHD-DR1.ACE2^Hu1^ in the lungs (Fig 4C). In the brain, we observed a tendency to a higher *hACE2* mRNA expression in HHD-DR1.ACE2^Hu3^ compared to HHD-DR1.ACE2^Hu2^. Although this trend did not reach significance, it was associated with a significantly higher permissiveness of HHD-DR1.ACE2^Hu3^ mice to Omicron BA.1 mice (Fig 4B).

Like the B6.K18-hACE2^IP-THV^ mice, the HHD-DR1.ACE2^Hu2^ and HHD-DR1.ACE2^Hu3^ mice were also strongly permissive to replication of Omicron BA.5 (Fig 5A). In contrast to the brains of HHD-DR1.ACE2^Hu2^ mice, which were not permissive to replication of Omicron BA.5, the brain of 3 out of 5 HHD-DR1.ACE2^Hu3^ mice had high cerebral Omicron BA.5 loads (Fig 5B).

**Figure 5.**
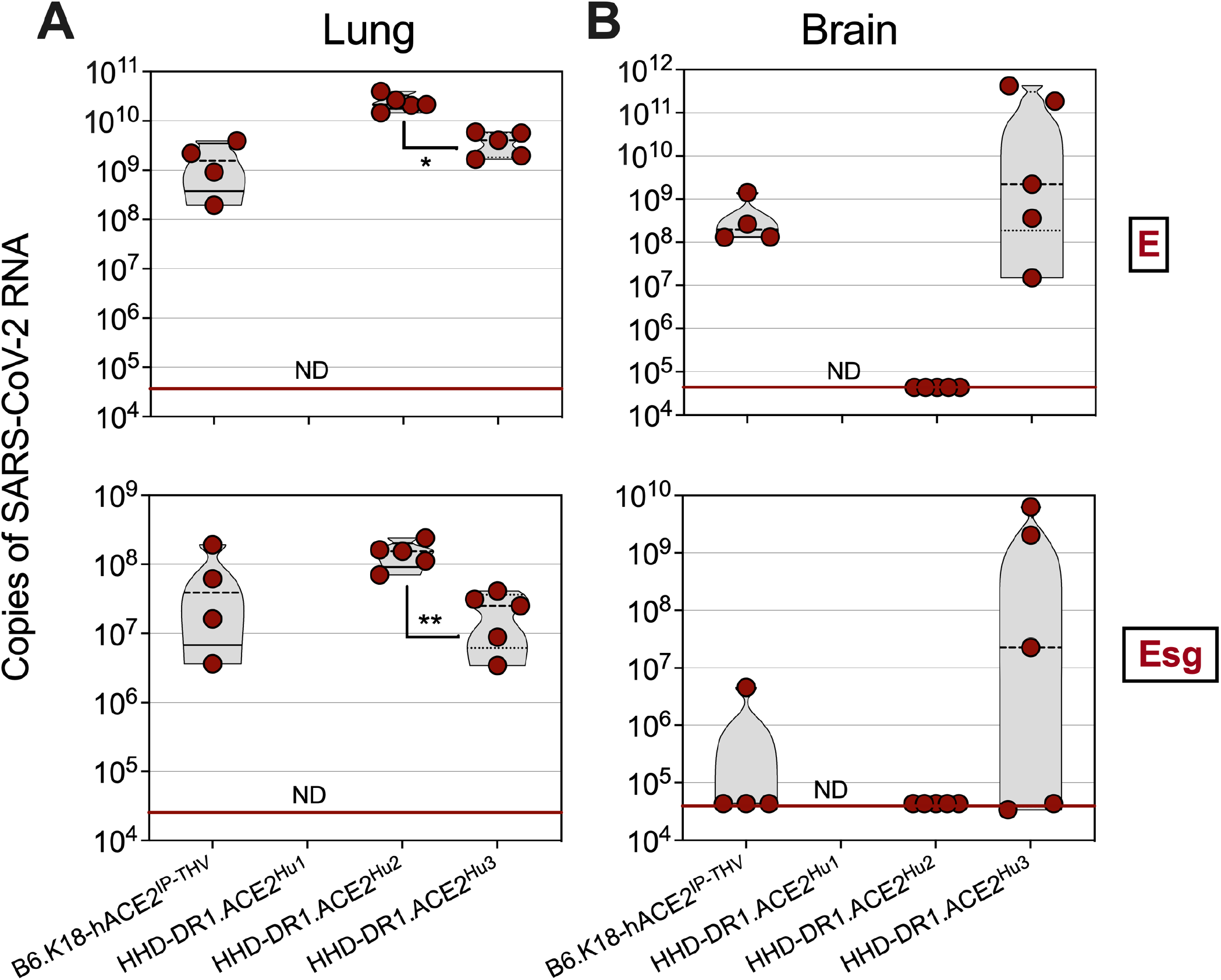
Permissiveness of HHD-DR1.ACE2Hu1/2/3 mice to replication of SARS-CoV-2 Omicron BA.5 sub-variant. HHD-DR1.ACE2Hu1/2/3 mice or B6.K18-hACE2IP-THV positive control mice were inoculated i.n. with 0.3 × 10^5^ TCID50/mouse of SARS-CoV-2 Omicron BA.5. **A-B** Lung **(A)** and brain **(B)** viral RNA contents were determined by E-specific (top) or sub-genomic Esg-specific (bottom) qRT-PCR at 4 dpi. Statistical significance was evaluated by Mann-Whitney test (*= *p* < 0.05, **= *p* < 0.01).

Therefore, permissiveness to the two Omicron sub-variants tested was observed in both lungs and brains of the MHC-humanized mice, reaching levels as high or even higher than in B6.K18-hACE2^IP-THV^

### Use of HHD-DR1.ACE2^Hu1^ to evaluate the efficacy of a lentiviral vector-based COVID-19 vaccine candidate

To conduct a vaccine efficacy evaluation in one of the HHD-DR1.ACE2^Hu^ strains, we first studied, in HHD-DR1 mice, the immunogenicity of the lentiviral vector-based COVID-19 vaccine candidate, namely “LV::S_Beta-2P_” (Ku *et al*., 2021a; Ku *et al*., 2021b; Majlessi & Charneau, 2021; Vesin *et al*., 2022). LV::S_Beta-2P_ encodes the full length sequence of Spike from the SARS-CoV-2 Beta variant, stabilized by K^986^P and V^987^P substitutions in the S2 domain (Vesin *et al*., 2022).

As we previously established that an intramuscular (i.m.) prime followed by an i.n. boost was the most efficacious scheme to achieve anti-SARS-CoV-2 protection, HHD-DR1 mice were primed (i.m.) with 1 × 10^8^ Transduction Units (TU) of LV::S_Beta-2P_ at wk 0 and then boosted (i.n.) with the same amount of this vaccine at wk 8. Control mice received an empty LV vector (Ctrl LV). The LV::S_Beta-2P_-immunized mice mounted high titers of serum anti-Spike IgG, albeit weaker than those detected in the conventional C57BL/6 mice (Fig S1) (see Discussion).

T-cell immunogenicity of LV::S_Beta-2P_ in HHD-DR1 mice was studied by IFN-γ ELISPOT epitope mapping of Spike, based on a 2D-peptide pool matrix (Hoffmeister *et al*, 2003), using splenocytes (Fig 6A, B). To do so, panels of 253 15-mers spanning the full-length Spike protein were organized into 32 pools, assigned to a 2D matrix in which each peptide was represented in two distinct pools. In this approach, the intersection of two immunogenic pools predicts presence of a potentially positive epitope. By analyzing 5 individual LV::S_Beta-2P_-vaccinated HHD-DR1 mice, we identified the positive IV, XII, XVI, XX, XXIII and XXVII peptide pools (Fig 6A, B). The intersection peptides, as well as S:511-525 (#103, VVLSFELLHAPATVC) and S:976-990 (#196, VLNDILSRLDKVEAE), previously identified in humans by others (Habel *et al*, 2020; Peng *et al*, 2020), were then individually tested in IFN-γ ELISPOT (Fig 6C). This assay clearly confirmed the immune-recognition of S:316-330 (#64, SNFRVQPTESIVRFP), S:511-525 (#103, VVLSFELLHAPATVC), S:536-555 (#108 and #109, NKCVNFNFNGLTGTGVLTES), S:816-830 (#164, SFIEDLLFNKVTLAD), and S:976-990 (#196, VLNDILSRLDKVEAE) by T cells from HHD-DR1 vaccinated mice. Interestingly, none of these identified T-cell epitopes overlapped with regions where mutations occurred in the Omicron Spike. This suggests that LV::S_Beta-2P_-induced T cells will recognize Omicron-infected host cells as efficiently as those infected with the ancestral virus.

**Figure 6.**
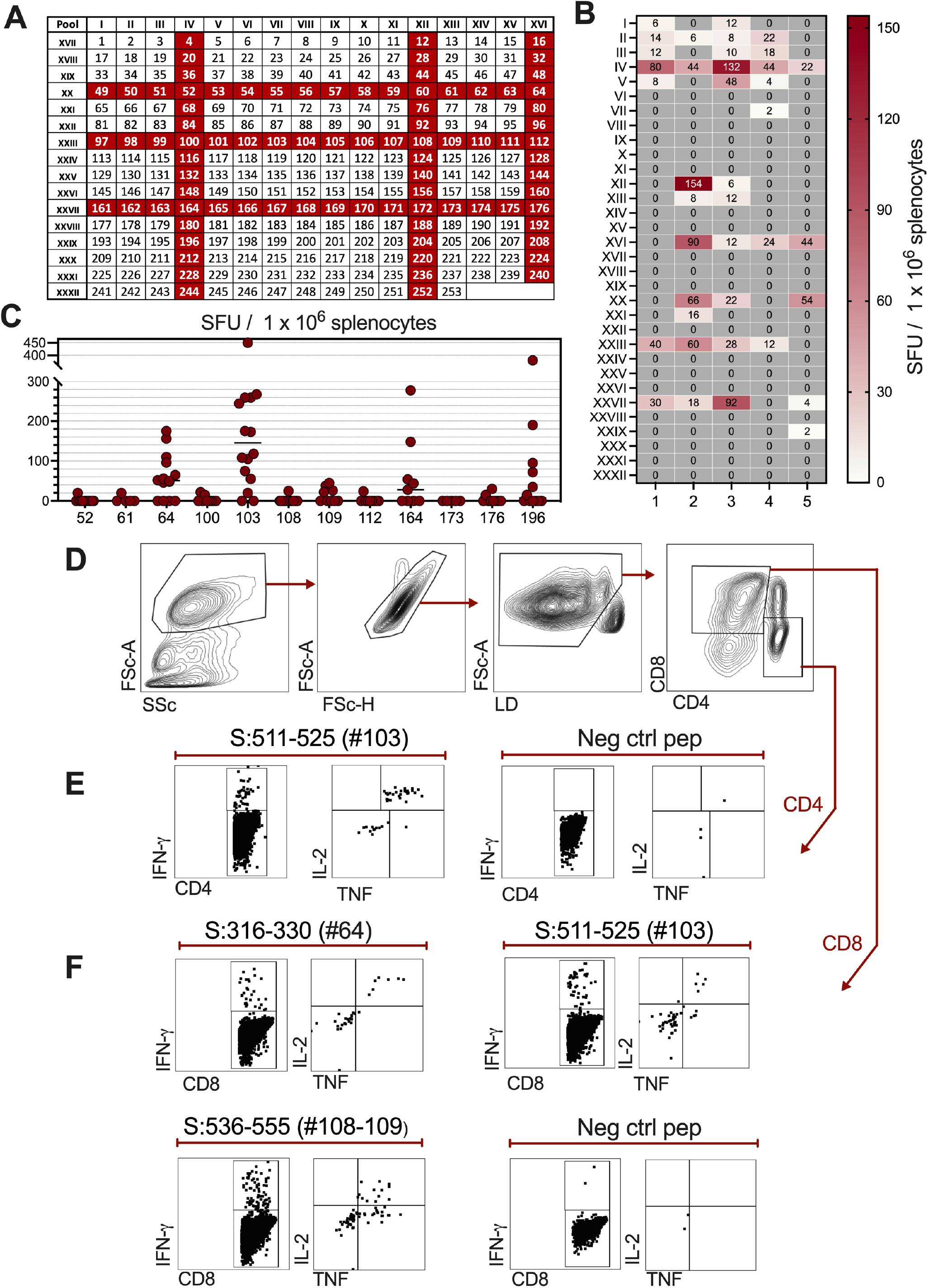
Spike T-cell epitope mapping in HHD-DR1 mice vaccinated with LV::SBeta-2P. HHD-DR1 mice were primed (i.m.) at wk 0 and boosted (i.n.) at wk 8 with a control empty lentiviral vector (Ctrl LV) or LV::SBeta-2P. **A** Panels of 253 15-mers spanning the Spike protein, were organized into 32 pools (I to XXXII) containing no more than 16 individual peptides. The peptide pools were assigned to a 2D matrix system in which each peptide was represented in two different pools. The intersection of two positive pools identified potentially positive peptides. **B** Splenocytes from HHD-DR1 mice (*n* = 5) immunized with LV::SBeta-2P or a Ctrl LV were harvested at wk 2 post boost and were analyzed in an IFN-γ ELISPOT after stimulation with each of the 32 distinct I to XXXII pools. Heat map representing the number of IFN-γ-producing cells per million of splenocytes responding to each peptide pool. For each peptide pool, the background generated by the splenocytes from Ctrl LV-injected mice was substracted. **C** Intersection or previously identified peptides were tested individually in ELISPOT. **D-F** The positive peptides identified were used in CD4 and CD8 surface staining and IFN-γ, IL-2, and TNF ICS and cytometric analysis to determine the T-subset specific to the corresponding epitopes. **(D)** Gating strategy. **(E)** CD4 and **(F)** CD8 T cells responding to the identified peptides.

Intracellular cytokine staining (ICS) and cytometric analysis performed with splenocytes from LV::S_Beta-2P_-vaccinated HHD-DR1 mice, stimulated in vitro with these identified peptides, showed that: (i) S:511-525 (#103, VVLSFELLHAPATVC) was recognized by CD4^+^ T cells and thus restricted by DRA01.01 + DRB1.01.01 MHC-II molecule (Fig 6D). In parallel, S:316-330 (#64, SNFRVQPTESIVRFP), S:511-525 (#103, VVLSFELLHAPATVC) and S:536-555 (#108 and #109, NKCVNFNFNGLTGTGVLTES) were recognized by CD8^+^ T cells and thus contain epitopes restricted by HLA 02.01 MHC-I molecule (Fig 6E). Note that S:511-525 (#103, VVLSFELLHAPATVC) contains both MHC-I- and -II-restricted epitopes. Other peptides detected positive in the ELISPOT (Fig 6C) were not identifiable by ICS, the most likely because of the weaker sensitivity of ICS compared to ELISPOT. Further analysis of these immunogenic regions by the SYFPEITHI software (http://www.syfpeithi.de/bin/MHCServer.dll/EpitopePrediction.htm) also indicated the presence of high-scoring predicted T-cell epitopes in all these identified regions **(Table 2S)**.

Then, HHD-DR1.ACE2^Hu1^ mice were primed i.m. at wk 0 and boosted i.n. at wk 3 with LV::S_Beta-2P_ or Ctrl LV, and were then challenged i.n. at wk 5 with 0.3 × 10^5^ TCID_50_ of SARS-CoV-2 Delta variant. LV::S_Beta-2P_ prime-boost vaccination of HHD-DR1.ACE2^Hu1^ mice led to statistically significant protection against SARS-CoV-2 replication in the lungs, as determined at 4 dpi (Fig 7A). At this time point, cytometric analysis of the lung innate immune cell subsets detected significantly lower percentages of CD11b^int^ NKp46^+^ Natural Killer (NK) cells and CD11b^+^ Ly6G^+^ neutrophils among the CD45^+^ cells in the vaccinated HHD-DR1.ACE2^Hu1^ mice, than in the control animals which received Ctrl LV (Fig 7B). NK and neutrophils have been associated with enhanced lung inflammation and poor COVID-19 outcome and their decrease is a biomarker of good prognosis in humans (Cavalcante-Silva *et al*, 2021; Masselli *et al*, 2020).

**Figure 7.**
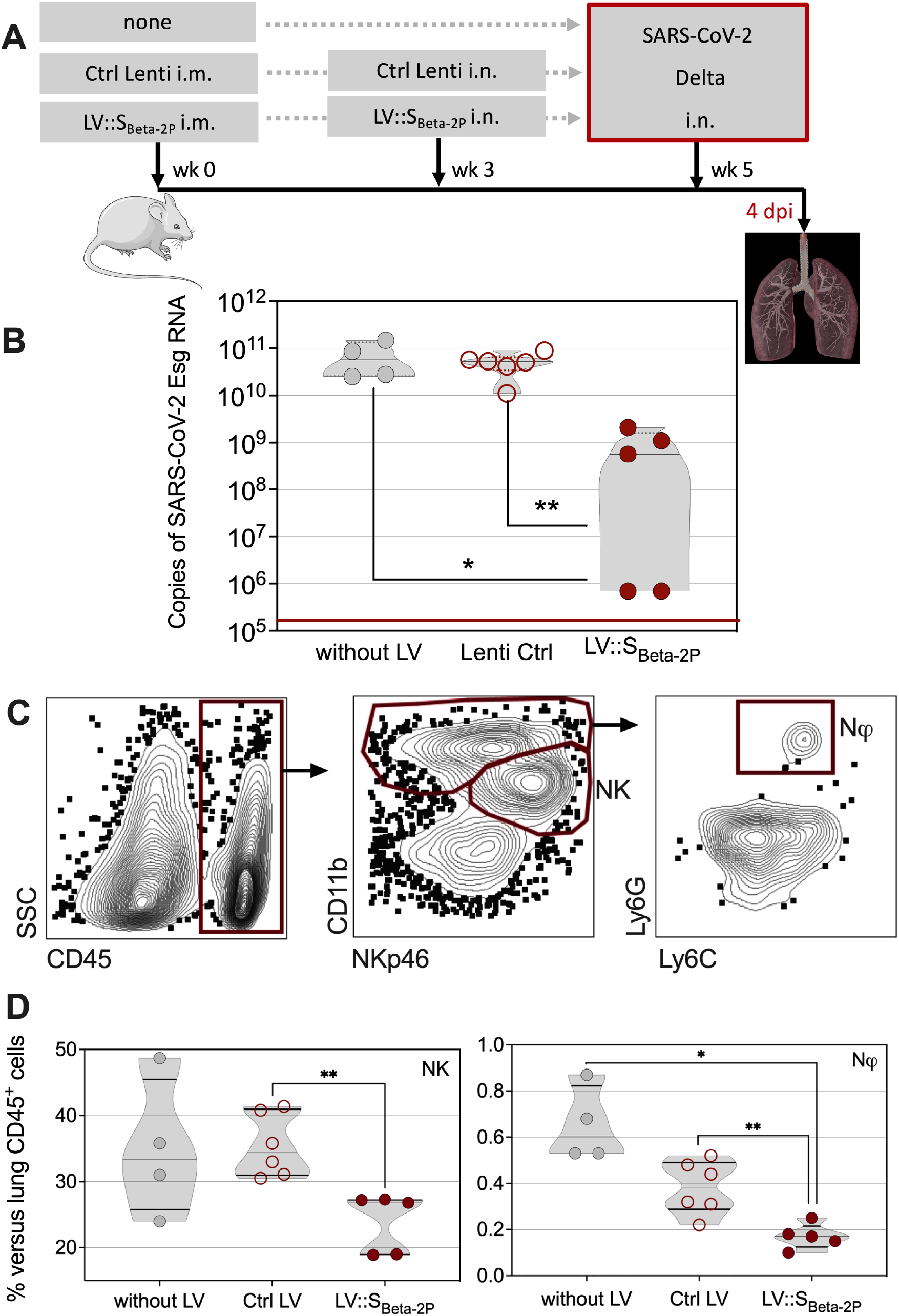
Use of HHD-DR1.ACE2^Hu1^ mice to evaluate the protective capacity of LV::SBeta-2P vaccine candidate against SARS-CoV-2. **A** Timeline of vaccination and SARS-CoV-2 challenge. **B** HHD-DR1.ACE2Hu1 mice were untreated or primed (i.m.) at wk 0 and boosted (i.n.) at wk 3 with 1 × 10^8^ TU/mouse of LV::SBeta-2P or Ctrl LV, before i.n. challenge at wk 5 with 0.3 × 10^5^ TCID50 of SARS-CoV-2 Delta variant. Lung viral Esg-specific RNA contents was evaluated by qRT-PCR at 4 dpi. Red lines indicate the detection limits. **C** Cytometric detection of neutrophils or NK cells in the lungs of untreated, Ctrl-Lenti-, or LV::SBeta-2P-vaccinated and challenged mice at 4 dpi. **D** Percentages of neutrophils or NK cells in the lungs of untreated, Ctrl LV-, or LV::SBeta-2P-vaccinated and challenged mice at 4 dpi. Percentages were calculated versus total lung live CD45+ cells. Statistical significance was evaluated by Mann-Whitney test (*= *p* < 0.05, **= *p* < 0.01).

Histological examination of SARS-CoV-2-inoculated mice pre-treated with Ctrl LV revealed a mild to moderate inflammation of the lung, with zones of interstitial infiltration sometimes accompanied by alveolar exudates (Fig 8A). Although the bronchiolar epithelium integrity was preserved, degenerative lesions of epithelium cells could be seen in the inflammatory areas. The histological scores are recapitulated in the heatmap (Fig 8B). In vaccinated HHD-DR1.ACE2^Hu1^ mice, the alveolo-interstitial syndrome was more limited in size, less severe (minimal to mild), and accompanied by discreet alterations of the bronchiolar epithelium.

**Figure 8.**
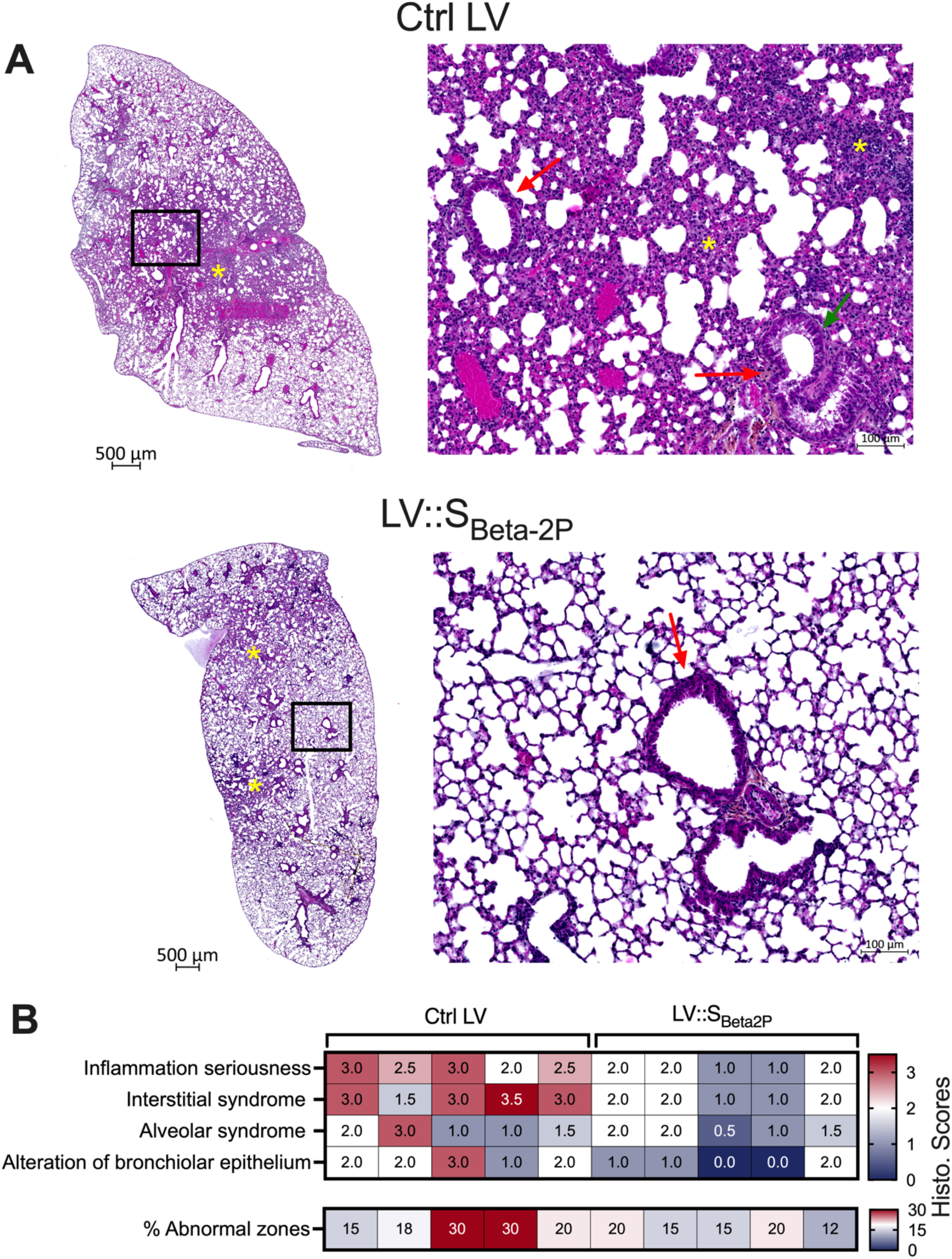
Lung histopathology of unvaccinated or LV::SBeta-2P-vaccinated HHD-DR1.ACE2^Hu1^ mice after inoculation with SARS-CoV-2. Mice are those detailed in the Figure 7. **A** Examples of haematoxylin-eosin saffron staining of whole-lung sections (left, scale bar: 500 μm) and close-up views (right, scale bar: 100 μm), at 4 dpi. Stars mark areas of minimal (bottom) or mild-to-moderate (top) infiltration. In the blow-up panels, red arrows indicate bronchiolar epithelia. While in the bottom blow-up the epithelium is subnormal, in the top panel epithelial degenerative lesions can be seen, e.g., perinuclear clear spaces (green arrow), associated with cells remnants and proteinaceous material in the bronchiolar lumen. Stars mark areas of minimal (bottom) or mild-to-moderate (top) infiltration. In the blow-up panels, red arrows indicate bronchiolar epithelia. While in the bottom blow-up the epithelium is subnormal, in the top panel epithelial degenerative lesions can be seen, e.g., perinuclear clear spaces, green arrow, and the lumen contains cells remnants and proteinaceous material. **B** Heatmap representing the histological scores for: (i) inflammation seriousness, (ii) interstitial syndrome, and (iii) alveolar syndrome, (iv) alteration of bronchial epithelium, and (v) % of abnormal lung zones.

Therefore, virological, immunological and histopathological criteria showed that HHD-DR1.ACE2^Hu^ mice can be used to evaluate protective potential of COVID-19 vaccine candidates.

## Discussion

We generated three new transgenic mice expressing: (i) hACE2 under the human K18 promoter, and (ii) only human – but not murine – MHC-I and -II molecules from HLA 02.01, DRA01.01, DRB1.01.01 alleles, widely expressed in human populations. The three strains, namely “HHD-DR1.ACE2^Hu1/2/3^”, are permissive to SARS-CoV-2 replication, yet each displays distinctive characteristics in terms of lung- and/or brain-specific permissiveness to viral replication, correlative with their site-specific *hACE2* mRNA transcription levels. Similar to B6.K18-hACE2^IP-THV^ strain (Ku *et al*., 2021a), and in clear contrast to B6.K18-ACE2^2Prlmn/JAX^ strain (Halfmann *et al*., 2022; McCray *et al*., 2007), one particularity of the HHD-DR1.ACE2^Hu1/2/3^ strains is their marked permissiveness to the replication of Omicron sub-variant(s). The high *hACE2* mRNA transcription in HHD-DR1.ACE2^Hu1/2/3^ strain, and notably in the HHD-DR1.ACE2^Hu3^ generated by lentiviral based transgenesis, is correlated with their permissiveness to SARS-CoV-2 Omicron replication. The reasons why the two independent lentiviral-based transgeneses, which gave rise to B6.K18-hACE2^IP-THV^ and HHD-DR1.ACE2^Hu3^ strains, led to better lung and still higher brain transgene expression are not clear. It is plausible that the lentiviral insertion sites in the host chromosome(s) lead to more accessible and more active transcription of the transgene. In addition, the Woodchuck hepatitis virus Posttranscriptional Regulatory Element (WPRE), present in the genetic material delivered by the lentiviral vector, enhances the transgene expression and transgenic mRNA export from the host nuclei (Zufferey *et al*, 1999), which is not the case for the transgene inserted by the conventional, DNA fragment-based transgenesis.

Among all SARS-CoV-2 variants of concern, Omicron harbors the highest number, i.e., 37 mutations in its Spike glycoprotein. Among these mutations, 15 are located inside the Receptor-Binding Domain (RBD). Inside the Omicron RBD, the 96 amino acid-long Receptor-Binding Motif (RBM) carries 10 mutations, i.e., N440K, G446S, S477N, T478K, E484A, Q493R, G496S, Q498R, N501Y and Y505H, among which, G446S, E484A, Q493R, G496S, Q498R and Y505H are unique to Omicron and are believed to reinforce by threefold the binding of its Spike to hACE2, compared to the Spike of ancestral or Delta SARS-CoV-2. This strengthened interaction occurs via increased electrostatic and hydrophobic interactions, and formation of ACE2 salt bridge and ACE2 hydrogen bond, improving the potential of Omicron to invade the host cells (Kumar *et al*, 2022; Suzuki *et al*, 2022; Zhao *et al*, 2022). On the other hand, the Omicron Spike protein is less efficiently cleaved into S1 and S2 subunits, reducing its fusiogenic potential and thereby its pathogenicity (Mlcochova *et al*, 2021; Saito *et al*, 2022). Other mutations elsewhere in the genome of Omicron may also be responsible its reduced pathogenic potential. Despite the increased affinity of Omicron RBM interaction with hACE2, replication of Omicron variants in the upper and lower respiratory airways of B6.K18-ACE2^2Prlmn/JAX^ transgenic mice is 3-to-5 log_10_ lower compared to the earlier SARS-CoV-2 variants (Halfmann *et al*., 2022). The increased permissiveness of HHD-DR1.ACE2^Hu1/2/3^ strains to Omicron, relative to that of the B6.K18-ACE2^2Prlmn/JAX^ mice (Halfmann *et al*., 2022), is probably largely explained by their higher expression of *hACE2* transgene transcription levels, because SARS-CoV-2 enters the host cells mainly via hACE2, although alternative routes of entry such as the TransMembrane Protease Serine 2 (TMPRSS2) or the cathepsin-dependent endocytic pathway have been reported (Balint *et al*, 2022). However, these are unlikely to be different in the B6.K18-ACE2^2Prlmn/JAX^, B6.K18-ACE2^IP-THV^ and HHD-DR1.ACE2^Hu1/2/3^ mouse strains, which harbor a common C57BL/6 genetic background.

As a first application of these strains, we demonstrated in HHD-DR1.ACE2^Hu1^ mice the efficacy of a lentiviral vector-based COVID-19 vaccine candidate. This vaccine induced anti-Spike serum IgG, CD4^+^ and CD8^+^ T cells (Ku *et al*., 2021a; Ku *et al*., 2021b; Majlessi & Charneau, 2021; Vesin *et al*., 2022). In the LV::S_Beta-2P_-immunized HHD-DR1 mice, the titers of anti-Spike IgG were lower than in their LV::S_Beta-2P_-immunized wild type C57BL/6 counterparts. This may be linked to the absence of free murine β2m in HHD-DR1 mice, which harbor only the human β2m covalently linked to the HLA 02.01. β2m contributes to the formation of “neonatal Fc receptor” (FcRn) for IgG (Zhu *et al*, 2002). In the absence of association with β2m, the FcRn heavy chain remains sequestered and unfunctional in endoplasmic reticulum. Increased clearance of IgG in β2m KO mice has suggested that FcRn protects IgG from degradation, and is thus critical in maintaining IgG levels in the circulation (Israel *et al*, 1996). In a non-mutually-exclusive manner, it is also possible that the size of the T-cell compartment in HHD-DR1 mice is reduced compared to wild-type mice. In this case, the CD4^+^ T-cell helper function may be attenuated, resulting in weaker antibody responses. Spike epitope mapping in LV::S_Beta-2P_-immunized HHD-DR1 mice allowed for the identification of both MHC-I- and II-restricted immunogenic Spike regions for humans. Furthermore, LV::S_Beta-2P_-vaccinated HHD-DR1^Hu1^ mice were protected from a challenge by SARS-CoV-2, as evidenced by virological, immunological and histopathological criteria, validating the HHD-DR1^Hu^ preclinical model for immunogenicity and vaccine evaluation investigations.

HHD-DR1.ACE2^Hu1/2/3^ murine strains provide new pre-clinical research tools for SARS-CoV-2 infection and research and development of new COVID-19 drugs or vaccines. These strains will notably pave the way for: (i) identification of SARS-CoV-2-derived epitopes recognized by human T cells, (ii) investigation of T-cell immunogenicity of new-generation COVID-19 vaccines, and (iii) evaluation of protective potential of new-generation COVID-19 vaccine, in small rodent models, in which the MHC-I and -II molecules are from widely expressed HLA alleles in human populations. In addition, since coronaviruses can infect the brain (Aghagoli *et al*, 2020; Ali Awan *et al*, 2021; Bourgonje *et al*, 2020), the HHD-DR1.ACE2^Hu2/3^ strains provide also valuable pre-clinical models for investigation of immune protection of the central nervous system against SARS-CoV-2.

## Material and Methods

### Mice

HHD-DR1 adult males and 3-week-old female mice were purchased from Charles River (Les Oncins, Saint Germain-Nuelles, France) and housed in ventilated cages under specific pathogen-free conditions at the Institut Pasteur animal facilities. B6CBAF1females were purchased from Janvier Labs (Janvier Labs, Le Genest-Saint-Isle). B6.K18-hACE2^IP-THV^ mice (Ku *et al*., 2021a; Vesin *et al*., 2022) were bred and housed in animal facilities of Institut Pasteur. All procedures were performed in accordance with the European and French guidelines (Directive 86/609/CEE and Decree 87-848 of 19 October 1987) subsequent to approval by the Institut Pasteur Safety, Animal Care and Use Committee, protocol agreement delivered by local ethical committee (CETEA #DAP20007, CETEA #DAP200058) and Ministry of High Education and Research APAFIS#31068-2021041613059523 v1, APAFIS#24627-2020031117362508 v1, APAFIS#28755-2020122110238379 v1.

### *hACE-2* transgenesis in HHD-DR1 mice

Zygotes from HHD-DR1 mice were microinjected with a 6.8-kb Hpal I-XbaI fragment from the pK18-hACE2 vector at 2 ng/μl (McCray *et al*., 2007), purchased from Addgene (Watertown, MA). Microinjected fertilized eggs were then transferred into pseudo-pregnant B6CBAF1 foster mothers (Janvier Labs). The progeny was studied for integration of *hACE2* gene by using hACE2-forward: 5’-TCC TAA CCA GCC CCC TGT T-3’ and hACE2-reverse: 5’-TGA CAA TGC CAA CCA CTA TCA CT-3’ primers in PCR, applied on genomic DNA prepared from toe clipping, as described previously (Ku *et al*., 2021a). PCR-identified *hACE2*^+^ founders were backcrossed to HHD-DR1 mice in order to preserve the state of homozygosity for HLA 02.01, DR_α_01.01, DR_β_01.01 humanized loci and for quintuple KO for β2m, H-2D^b^, I-A_α_^b^, I-A_β_^b^ and I-E_α_^b^ murine loci.

It is noteworthy that the female founder F0#9 gave birth to only one *hACE2*^+^ male, in the progeny of which only females were *hACE2*^+^, indicating that the *hACE2* transgene is on the X chromosome. Therefore, in the resulted strain, the *hACE2* transgene insertion is not lethal since males are in good state. The transition to homozygosity will be easy as it can be determined by a simple progeny test with which a *hACE2*^+^ male is “pseudo homozygous”, and if crossed with heterozygote females, will give rise to 50% of homozygote females. In addition, whether male or female, the founder F0#9 descendance will express only one functional *hACE2* allele. In fact, in females this will result from methylation of the *hACE2* allele on the second X chromosome.

### SARS-CoV-2 inoculation and measurement of viral RNA contents

The *hACE2*^*+*^ offspring were inoculated i.n., with 0.3 × 10^5^ TCID_50_ – contained in 20 μl – of the Delta/2021/I7.2 200 (Planas *et al*., 2021), Omicron BA.1 (Planas *et al*, 2022) or BA.5 SARS-CoV-2 clinical isolate. For i.n. inoculation, mice were anesthetized by i.p. injection of Ketamine (Imalgene, 80 mg/kg) and Xylazine (Rompun, 5 mg/kg). Animals were then housed in an isolator in BioSafety Level 3 animal facilities of Institut Pasteur. Viral RNA contents were determined in the organs by quantification of total or sub-genomic RNA from E_CoV-2_ by quantitative real-time (qRT)-PCR. The Esg RT-PCR measures only active viral replication (Chandrashekar *et al*, 2020a; Ku *et al*., 2021b; Tostanoski *et al*., 2020; Wolfel *et al*., 2020).

### qRT-PCR for Inflammation study

The qRT-PCR quantification of inflammatory mediators in the lungs and brain was performed as detailed elsewhere (Ku *et al*., 2021b) on total RNA extracted by TRIzol reagent (Invitrogen) and stored at −80°C. The RNA quality was first assessed using a Bioanalyzer 2100 (Agilent Technologies). The RNA Integrity Number (RIN) was 7.5 — 10.0. RNA samples were quantitated using a NanoDrop Spectrophotometer (Thermo Scientific NanoDrop). One μg of each RNA was used per qRT-PCR sample in 96 wells plate.

### ELISPOT assay and T-cell epitope mapping

Splenocytes from individual mice were homogenized and filtered through 70 μm-pore filters and centrifuged at 300g for 10 min. Cells were treated with Red Blood Cell Lysing Buffer (Sigma) and resuspended at 2.5 × 10^7^ cells/mL in complete α-MEM medium containing 10% heat-inactivated FBS, 100 U/mL penicillin and 100 μg/ml streptomycin, 1 × 10^−4^ M non-essential amino-acids, 10 mM Hepes, 1 mM sodium pyruvate and 5 × 10^−5^ M of β-mercapto-ethanol. For each mouse, 100 μl of cell suspension (2.5 × 10^5^ splenocyte/well) were stimulated overnight (∼16h) with 100 μl of SARS-CoV-2 matrix peptide pools (2 μg/ml of each peptide, Mimotopes, UK), SARS-CoV-2 individual peptides (2 μg/ml) or blank control (complete medium with DMSO) onto sterile nitrocellulose MSIP 96-well plates (Millipore, Bedford, MA) coated with capture antibodies against mouse IFN-γ. Panels of 253 15-mer peptides spanning the prototype Spike were organized into 32 pools. The overlapping peptide pools were assigned to a 2D matrix in which each peptide was represented in 2 different peptide pools. In this way, the intersection of two positive pools identifies a potentially positive peptide (Hoffmeister *et al*., 2003). Fifteen-mer candidate peptides at the intersections of the positive matrix-pools were tested individually for confirmation. The results were expressed as IFN-γ spot forming units (SFU) per 10^6^ splenocytes and were determined using on an ELISpot analyzer (ImmunoSpot^®^, CTL, Germany). To quantify antigen-specific responses, mean spots of the control wells were subtracted from the positive wells.

### Intracellular cytokine staining

Splenocytes were plated at 4 × 10^6^ cells/well in 24-well plates and co-cultured during 6h in the presence of 10 μg/ml of appropriate peptide, 1 μg/ml of anti-CD28 (clone 37.51) and 1 μg/ml of anti-CD49d (clone 9C10-MFR4.B) mAbs (BD Biosciences). During the last 3h of incubation, a mixture of Golgi Plug and Golgi Stop (BD Biosciences) were added. Cells were then collected, washed and stained for 25 min at 4°C with a mixture of Near IR Live/Dead (Invitrogen), FcγII/III receptor blocking anti-CD16/CD32 (clone 2.4G2), PerCP-Cy5.5-anti-CD3ε (clone 145-2C11), PE-Cy7-anti-CD4 (clone RM4-5) and BV711-anti-CD8 (clone 53-6.7) mAbs (BD Biosciences or eBioscience). Cells were washed in FACS buffer, permeabilized in Cytofix/Cytoperm kit (BD Bioscience), washed twice with PermWash 1X buffer and incubated with a mixture of BV421-anti-IL-2 (clone JES6-5H4), FITC-anti-TNF (MP6-XT22), and APC-anti-IFN-γ (clone XMG1.2) mAbs (BD Biosciences), during 25 min at 4°C. Cells were then washed in PermWash and then in FACS buffer and fixed with Cytofix (BD Biosciences) overnight at 4°C. Samples were acquired in an Attune NxT cytometer system (Invitrogen) and data were analyzed using FlowJo software (Treestar, OR, USA).

### Vaccine assay

Methods used for the measurement of Spike-specific antibody and T-cell responses were described elsewhere (Ku *et al*., 2021b). Mice were immunized (i.m. or i.n.) with 1 × 10^8^ TU of LV::S_Beta-2P_, contained in 50 μl for i.m. injection and in 20 μl for i.n injection. Mice were inoculated i.n. with 0.3 × 10^5^ TCID_50_ of the Delta variant of SARS-CoV-2 clinical isolate and housed in filtered cages in an isolator in biosafety level 3 animal facilities. The organs recovered from the infected animals were manipulated according to the approved standard procedures of these facilities in a BSL3 laboratory.

### Lung histopathology

Histopathological study was performed on the left lung lobes, fixed in formalin for 7 days, then transferred in ethanol and embedded in paraffin. Five μm-thick sections were stained with hematoxylin and eosin. Lesions were scored on images acquired in a double-blinded manner on an Axioscan Z1 Zeiss slide scanner, using the Zen 2 blue edition software.

### Statistical analyses

Statistical significances were determined by use of two-tailed unpaired t test. When indicated data were subjected to Pearson correlation coefficient analyses. All statistical analyses were performed by use of Prism GraphPad v9.

## Data Availability

The published article includes all datasets generated and analyzed during this study. All plasmids, LV and mouse strains generated in this study will be available under MTA for research use. Further information and requests for resources and reagents should be directed to and will be fulfilled by the corresponding author Laleh Majlessi.

## Acknowledgments

The authors are grateful to Magali Tichit and Johan Bedel for excellent technical assistance in preparing histological sections. The SARS-CoV2 variant Delta/2021/I7.2 200 was supplied by the Virus and Immunity Unit (Institut Pasteur, Paris, France) headed by Dr. Olivier Schwartz. The SARS-CoV-2 Omicron BA.1 variant was initially supplied by the Virus and Immunity Unit (Institut Pasteur, Paris, France) headed by Dr. Olivier Schwartz, and was provided to our lab by Matthieu Prot and Etienne Simon-Lorière (G5 Evolutionary Genomics of RNA Viruses, Institut Pasteur, Paris, France). The strain hCoV-19/France/BRE-IPP34319/2022 (Omicron BA.5) was supplied by the National Reference Centre for Respiratory Viruses hosted by Institut Pasteur (Paris, France) and headed by Pr. Sylvie van der Werf. The human sample from which strain hCoV-19/France/BRE-IPP34319/2022 was isolated have been provided by Dr; Marque-Juillet from Laboratoire Alliance Anabio, in Melesse.

This work was supported by Institut Pasteur and TheraVectys.

## Author Contributions

Funding: MB, PC, Study concept and design: MB, PC, FL, FLV, LM, acquisition of data: FLC, PA, MB, BV, IF, KN, LM, lung histopathology: FG, DH, transgenesis experiments and breeding: SC, DC, YS, FLV, construction and production of lentiviral vector for transgenesis: FM, AN, CB, FA, analysis and interpretation of data: PC, FL, FLV, LM, drafting of the manuscript: LM.

