## Supplemental Information for "Mice Humanized for Major Histocompatibility Complex and Angiotensin-Converting Enzyme 2 with High Permissiveness to SARS-CoV-2 Omicron Replication"

**Table S1. Main features of the HHD-DR1$^{Hu}$ strains**

| **Transgenesis approach** | **Vector** | ***hACE2*$^{+}$ founders (sex)** | **Name** | **Permissiveness to SARS-CoV-2 Delta variant** | | **Permissiveness to SARS-CoV-2 Omicron BA.1** | | **Permissiveness to SARS-CoV-2 Omicron BA.5** | |
|---|---|---|---|---|---|---|---|---|---|
| **Micro-injection into fertilized HHD-DR1 eggs** | | | | **Lung** | **Brain** | **Lung** | **Brain** | **Lung** | **Brain** |
| Pronuclear DNA | pK18-hACE2 DNA | F0#8 (male) | — | - | - | - | - | - | - |
| | | F0#9 (female) | HHD-DR1.ACE2$^{Hu1}$ | + | - | + | ND | ND | ND |
| | | F0#11 (female) | HHD-DR1.ACE2$^{Hu2}$ | + | + | + | - | + | - |
| Sub-zonal under pellucida | LV::pK18-hACE2 | F30# (male) | — | - | - | - | - | - | - |
| | | F0#47 (male) | HHD-DR1.ACE2$^{Hu3}$ | + | + | + | + | + | +/- |

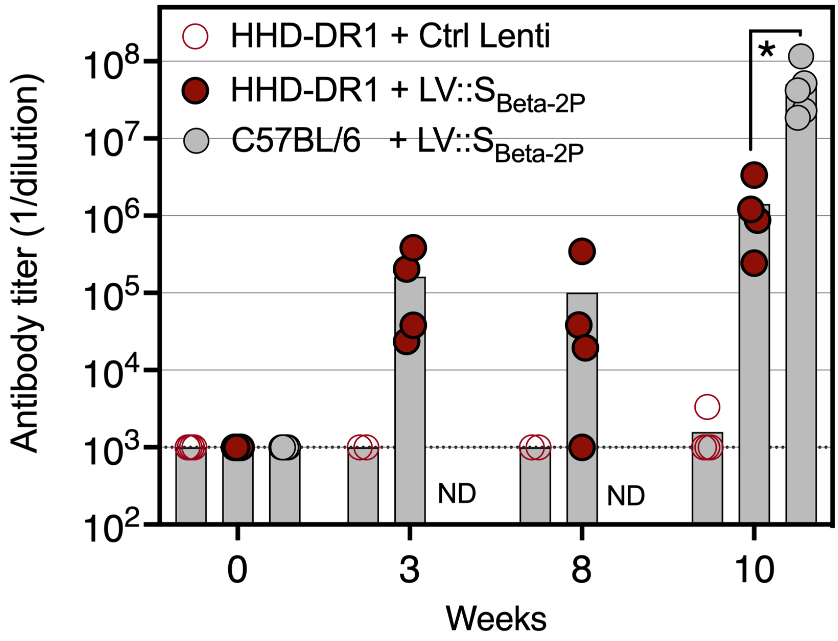


**Figure S1. Anti-Spike antibody response of HHD-DR1 mice to LV::S$_{Beta-2P}$ vaccination. (A)** HHD-DR1 mice were primed (i.m.) at wk 0 and boosted (i.n.) at wk 8 with a control empty lentiviral vector (Ctrl LV) or LV::S$_{Beta-2P}$. Anti-Spike IgG titers were studied in the sera before and after the boost. The antibody titers shown for wk 8 those determined in the sera before the i.n. boost. At wk 10, sera from C57BL/6 wild type mice vaccinated with LV::S$_{Beta-2P}$ were also studied. (ND = Not determined). Statistical significance was determined by Mann-Whitney test (*= $p < 0.05$).

**Table S2. Spike-derived T-cell epitopes identified by epitope mapping in HHD-DR1 mice.**

| Immunogenic regions identified in HHD-DR1 mice | a.a. sequence | Restricting element | «SYFPEITHI*» Score |
|---|---|---|---|
| S::316-330 (#64) | SNFR**V**QPT<u>E</u>SI**V**RFP | HLA 02.01 | 16 |
| S:511–525 (#103) | VVLSFE**L**LHA<u>P</u>AT**V**C<br>VV**L**SFE<u>L</u>LH**A**PATVC | | 23<br>19 |
| S:536-555 (#108 and #109) | NKCVNFNFNG**L**TGT<u>G</u>VL**T**ES | | 18 |
| S:816-830 (#164) | SFIED**L**LFN<u>K</u>VT**L**AD<br>SFIEDL**L**FNK<u>V</u>TL**A**D | | 24<br>22 |
| S:976-990 (#196) | V**L**NDI<u>L</u>SR**L**DKVEAE | | 27 |
| S:511–525 (#103) | **V**VL**S**F**E**LL**H**APATVC | DRA01.01 + DRB1.01.01 | 16 |

*http://www.syfpeithi.de/bin/MHCServer.dll/EpitopePrediction.htm: This algorithm predicts the ligation strength to MHC molecules. The score is defined by the probability for an a.a. sequence of being processed and presented by various MHC restricting elements. The highlighted 9-mers represent the potential MHC-I-restricted epitopes. The anchor residues are indicated in bold and the auxiliary anchors are underlined.
